# Prosociality in a despotic society

**DOI:** 10.1101/2022.08.07.503078

**Authors:** Debottam Bhattacharjee, Eythan Cousin, Lena S. Pflüger, Jorg J.M. Massen

## Abstract

Humans possess remarkable prosocial tendencies beyond the confinement of kinship, which may be instrumental in promoting cooperative interactions and sociality at large. Yet, prosociality is an evolutionary conundrum as it does not provide immediate benefits to the actor. The ‘domestication’ and ‘cooperative-breeding’ hypotheses postulated that enhanced social tolerance and inter-individual dependence could nonetheless facilitate the evolution of prosociality. However, inconsistent results due to varying experimental paradigms, and restricted focus of research on tolerant and cooperatively breeding species, have impeded our understanding so far. Albeit counterintuitively, despotic societies with relatively low social tolerance levels represent an excellent opportunity to investigate prosociality due to their kin favoritism and potential interdependence among individuals in terms of social support. Japanese macaques (*Macaca fuscata*) live in strictly hierarchical matrilineal societies, where kin members have strong social bonds. Additionally, support from non-kins can be crucial to form coalitions and rank up in the hierarchy. Using a group-service paradigm, we tested prosociality in a semi-free-ranging group of Japanese macaques. In contrast to currently existing evidence, we found that individuals (n=25) can act prosocially and at comparably high rates as cooperative breeding- or self-domesticated species. The macaques benefitted not only their kin members but other individuals to whom they showed relatively high social tolerance. We emphasize the roles of complex socio-ecological conditions in facilitating individual prosocial tendencies. Furthermore, these results call for a novel evolutionary framework regarding prosociality that focuses on different forms of interdependence and expands beyond cooperative breeding- and (self-)domesticated species.

**Significance statement:** What made humans so incredibly prosocial? Examining the evolutionary trajectory of prosocial acts led comparative psychologists to explore various taxa. Empirical evidence so far suggests that enhanced social tolerance and interdependence among individuals facilitate prosociality. Conventionally, despotism is characterized by low group-level tolerance, yet kin favoritism, nepotism, and high interdependence (in support and coalition formation) are also fundamental properties of despotic societies. Under such complex socio-ecological conditions, individual prosocial acts could thus be vital. We found, for the first time, high levels of prosociality in the very despotic Japanese macaques. Individuals benefitted both kin-relatives and others to whom they showed relatively high dyadic social tolerance. This study signifies that prosociality can be favored even in a highly despotic society.

## Introduction

Prosociality, the tendency to benefit others at low or no cost, is ubiquitous in human societies and is proposed to be a significant driver of the evolution of cooperative interactions and broadly sociality (1, 2). Yet enacting such behavior leads to no direct or immediate gain for the actor, making prosociality an evolutionary puzzle. In search of the evolutionary trajectory of prosociality, comparative psychologists examined and indeed found evidence of prosocial tendencies in a range of species other than humans (3, 4). While explaining the emergence and maintenance of prosociality remains challenging, two different hypotheses have gained empirical support, both highlighting the importance of enhanced social tolerance among group members. The self-domestication hypothesis considers prosociality a by-product of selection against reactive aggression, resulting in increased social tolerance (5, 6). On the other hand, the cooperative breeding hypothesis specifically points out allomaternal care, in other words, considerable inter-individual dependence in a group (7), to be a driving force facilitating prosociality (3, 5, 8–10). The two hypotheses are not mutually exclusive, and their proposed explanatory mechanisms seem to work both proximate and ultimate, thus at two different levels. Yet, variations in inter-individual prosocial motivations, even among cooperatively breeding or self-domesticated species, are poorly understood. Therefore, it is imperative to ascertain what underlying mechanisms predict prosociality at the group *and* individual levels and their context specificities.

The potential extent of interdependence is ample and may be predicted by factors like kin relationships and social support, leading to varying degrees of tolerance among members even within a social group, which may, in turn, lead to different levels of prosociality. Despotic societies are generally characterized by low group-level social tolerance and high dominance asymmetries (11, 12). Yet despotism favors kin, i.e., genetically closely related individuals may gain adequate benefit out of nepotistic acts (11). Also, social support and coalition formation are paramount in these societies to increase rank in the hierarchical system (13). While despotic societies are “conventionally” not expected to behave prosocially (11), from a ‘Machiavellian Intelligence’ (14, 15) perspective, prosociality, restricted to certain partners and contexts, could be expected; there also is some empirical evidence for prosociality in relatively despotic long-tailed macaques (16), capuchin monkeys (17), and chimpanzees (18–20), but see (21–23). These results, however, were obtained using different set-ups and procedures compared to those which found evidence of prosociality for the cooperative breeding and self-domestication hypotheses (3,5,9,10). Hence, testing more despotic species using similar set-ups and procedures is paramount for making clear inferences. Nevertheless, although these findings indicate a nepotistic influence on prosociality, prosocial attitudes toward non-kin members were also observed. Furthermore, these authors stress examining multiple groups due to evident inter-group variability in prosociality; and investigating species that show even steeper dominance hierarchies as there is a great lack of prosociality studies in despotic species.

Japanese macaques (*Macaca fuscata*) are a highly despotic species with remarkably low group-level social tolerance and high dominance asymmetry (11, 24). Yet, they exhibit strong matrilineal bonds with clear signs of nepotistic acts (25). Moreover, coalition formation (26) or alternate social strategies have been found to increase individual benefits within the hierarchy (e.g., males coalitions: (27)); or females originating from large matrilines competing with their sisters and therefore form non-kin alliances with higher-ranked individuals to “socially” outrank their higher ranked sisters (28). Japanese macaques are also capable of cooperative problem-solving in experimental settings in both semi-free and wild environments (29, 30), for which prosociality is considered to be a pre-requisite (31). Yet, previous studies found no evidence of prosocial tendencies in Japanese macaques in captivity (3, 10). The current study uses a well-established group service food provision paradigm (3, 5, 9, 10, 32) (see **Figure 1**) to explore prosociality in semi-free-ranging Japanese macaques. Our research offers several advantages over the previous studies on (Japanese macaque) prosociality – (a) a semi-free-ranging group with naturalistic conditions, i.e., high ecological validity, (b) a relatively large sample size; i.e., a group of over 170 macaques, out of which 25 participated in our experiment voluntarily (**Table S1**), (c) individual-level representation of prosocial motivations (if any), and (d) both group level- as dyadic social tolerance measures. We hypothesize that the complexity of a large despotic society creates interdependencies among its members favoring prosociality, and that close kin members (i.e., matrilines) display prosocial tendencies restricted to each other and more than non-kin. We also hypothesize that high social tolerance measures between non-kin dyads would positively predict the likelihood and magnitude of prosociality.

**Figure 1.**
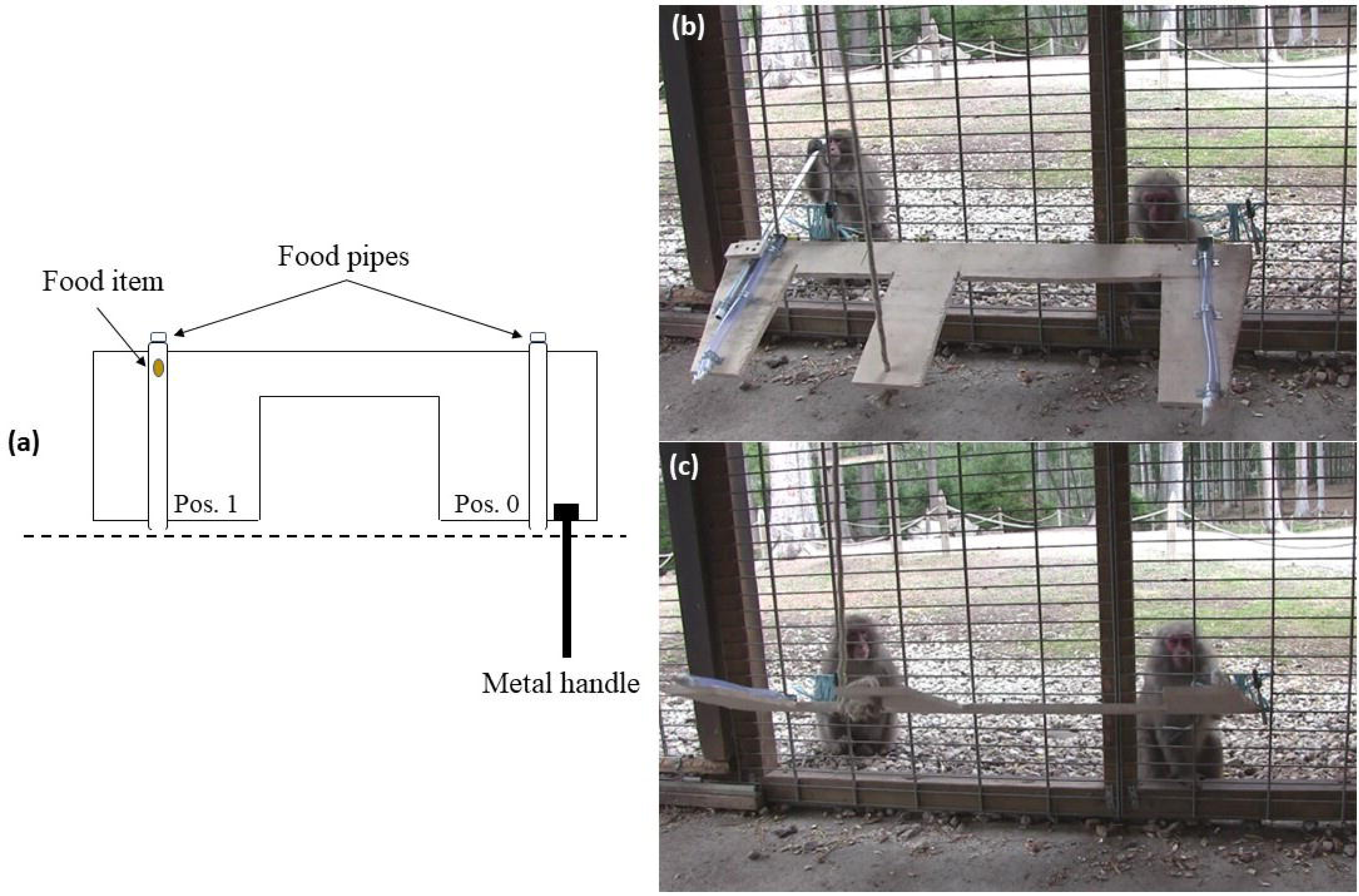
Experimental set-up of the group service paradigm. **(a)** In this seesaw mechanism, a wooden board (length ∼ 1.5 m) had two transparent plastic pipes (Ø ∼ 3 inches) attached to two extremes (Pos. 0 or providing end and Pos. 1 or receiving end) through which food rewards (i.e., peanuts) could move. By default, the board was tilted towards the experimenter present inside the research hut (step **(b)** in the figure). A metal handle was connected to the board (next to Pos. 0) and projected outside, which upon pressing, could tilt the board towards the macaques (step **(c)**), resulting in food rolling down through the pipe(s) (if present) and coming in reach for individuals. Only an attempt to press, i.e., the release of the handle halfway, caused the seesaw to move back to its original position, and eventually, food (if present) would roll out of reach of the individuals again. Therefore, it was essential to fully press and hold the handle to get the food rewards. Notably, an individual pressing and holding the handle could not reach Pos. 1 due to the length of the board. **(b)** and **(c)** The consecutive steps show how an adult male Japanese macaque is providing food to an unrelated adult female.

## Results

### Group and dyadic social tolerance

The group-level social tolerance was measured by looking at the evenness of food distribution. Phase 2 of the group service paradigm was explicitly designed to do so (see **Materials and Methods, SI Movie 1**). We calculated Pielou’s *J*′ (9, 10, 33) and obtained a value of 0.13. This suggests a highly uneven distribution of food (**Figure S1**) and, thus, a low group-level social tolerance (3, 5).

We conducted a string pulling task (see **Materials and Methods, SI Movie 2**) where individuals could obtain food simultaneously or monopolize it. The strength of dyadic social tolerance was defined by the instances where two individuals retrieved food rewards simultaneously. We found a dyadic social tolerance range from 0 to 18 (see **Supplementary Information** for details).

### Group and individual prosocial level presses

Individuals were presented with the opportunity to learn the contingencies of the procedure over five consecutive sessions per condition. Based on responses from the final two sessions (cf. 3, 5, 9, 10, 29), we observed a significant difference in terms of pressing the handle across different experimental conditions. Individuals pressed the handle to provide food to their conspecifics (**SI Movie 3**) significantly more often in the test (88%), where food was in turn also delivered to a conspecific, than in the empty- (36%, GLMM: z = -8.460, p < 0.001, **SI Movie 4**) and blocked control (35%, GLMM: z = -8.680, p < 0.001, **SI Movie 5**) conditions (**Figure 2**), where there was either no food or where the access to the food for the recipient was blocked, respectively. As the blocked control sessions were conducted after both the test and empty-control sessions (alternating with each other), we conducted two re-test and re-empty control sessions (again alternately) to account for order effects. During re-test sessions, the percentage of press (test = 85.2%, empty control = 30%) was comparable to earlier sessions (test = GLMM: z = 1.167, p = 0.24; empty control: GLMM: z = 0.511, p = 0.61; **Figure 3**), indicating no bias due to the pre-determined order of the experimental phases.

**Figure 2.**
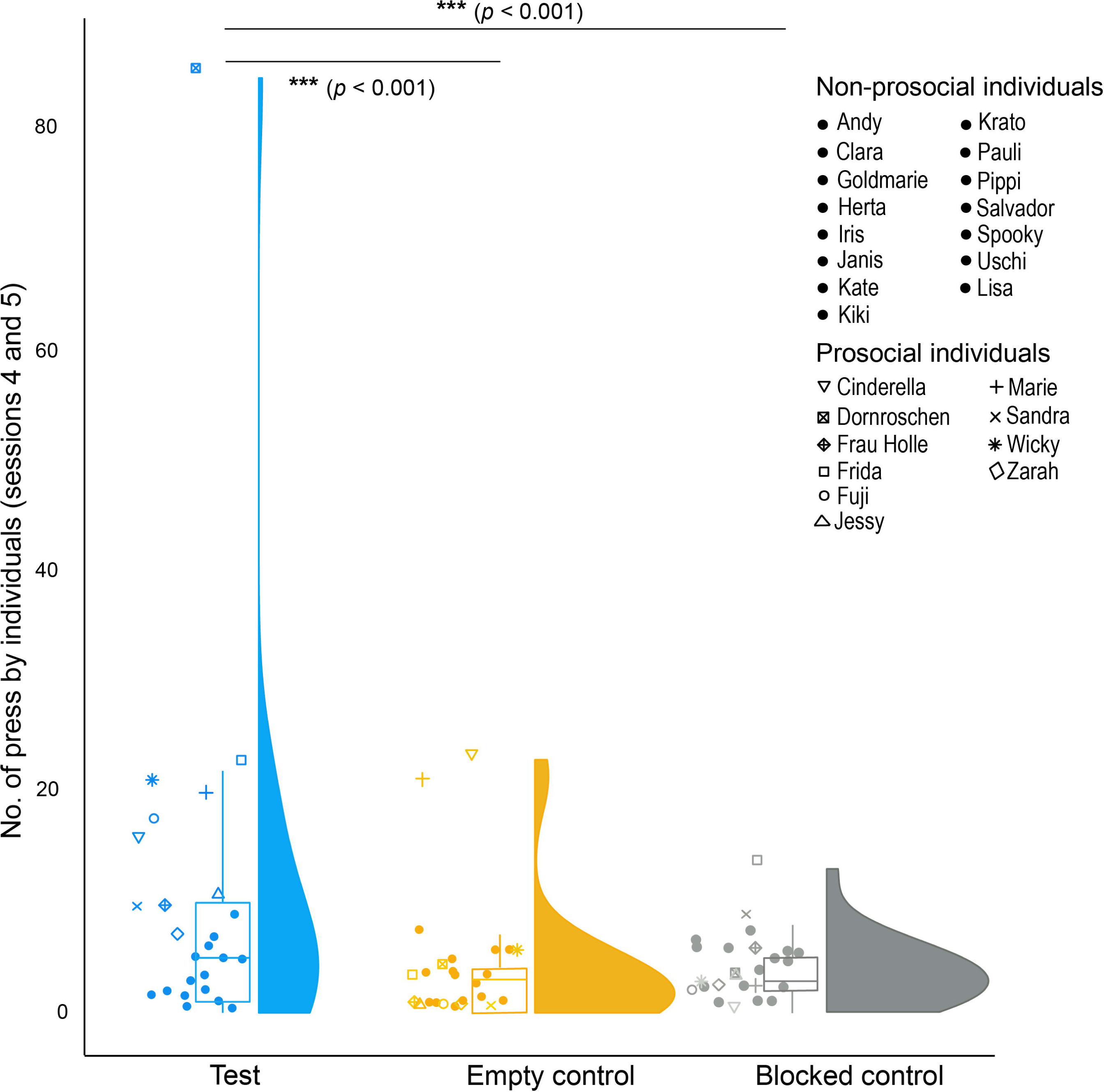
Overview of the number of ‘prosocial’ presses across different experimental conditions. Half-violin plots indicate the distribution of the data. Solid dots indicate the raw values of the non-prosocial individuals, whereas different symbols denote the raw values of the prosocial individuals (see identities in the figure). The boxes illustrate the interquartile range, horizontal bars inside the boxes indicate median values and whiskers indicate the range of the data, excluding outliers.

**Figure 3.**
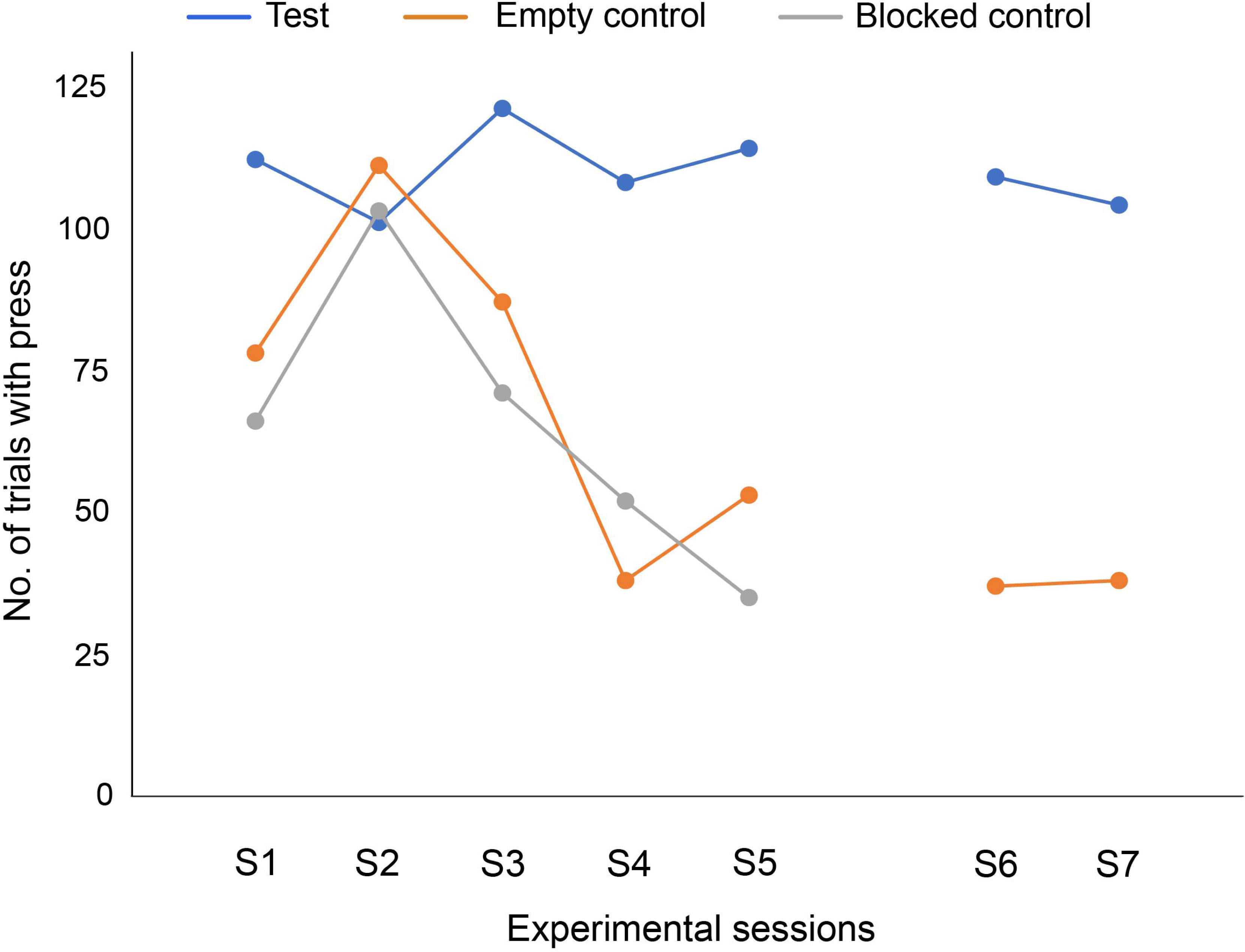
Trials with presses across sessions and experimental conditions. S1-S5 are regular sessions, while S6 and S7 indicate re-test and re-empty control sessions. The number of presses was comparable between regular test and re-test (85.2%), and regular empty control and re-empty control (30%) sessions.

Moreover, a sustained rate of pressing the handle across all test sessions was observed, this in contrast to the control sessions (**Figure 3**), confirming that the animals first had to learn the contingencies of these controls, whereas their prosocial tendencies seemed genuine.

At the individual level, we found that three macaques pressed the handle significantly more often in the test than in both control conditions, whereas seven of them pressed significantly more often in the test than in at least one of the control conditions. We considered these ten individuals to have “prosocial tendencies” (henceforth “prosocial” individuals) compared to others that we considered to have none (**Figure 2, Table S2**).

### Latency to press the handle

We found the latencies of pressing the handle to be different across the three conditions (**Figure S2**). Individuals pressed the handle faster in the test condition (18.32 ± 22.31 sec) as compared to both empty (45.37 ± 29.18 sec, LM: t = 9.584, p < 0.001) and blocked (44.52 ± 32.18 sec, LM: t = 9.047, p < 0.001) control conditions, again suggesting that the presses in the test condition were genuine and those of the control conditions needed more elaboration. Note that here we only considered trials when there was a press; therefore, trials without a press were discarded from this analysis.

### Food provision

Overall, food provisions, i.e. prosocial presses that also led to a recipient getting the food, occurred in 72.8% of the trials by all the individuals. For a more conservative measure of prosocial provisioning, we corrected this by using only the prosocial individuals who seemed to understand the consequences of their actions in the group-service paradigm, and the value was found to be 68.8%. Notably, even though it pressed more often in the test than in empty control, one individual did not provide any food to others. Thus, we excluded her from the list of prosocial individuals due to a lack of clear evidence of prosocial provisioning.

Although the prosocial individuals seemed to also provision food at a sustained rate (**Figure S3**), there was a significant difference between session one (no. of provisions ± SD: 3.9 ± 9.02) and five (8.8 ± 9.67) of the test condition (GLMM: z = 2.478, p = 0.013), indicating an increase in food provision from session one to five. Given the sustained rates of pressing by the focal subjects/prosocial individuals across sessions, these results suggest better coordination between actors and recipients.

Furthermore, for prosocial individuals, we compared the number of food provisions and food received (**Table S3**). These individuals provided food to others (no. of provisions ± SD: 19.11 ± 22.83) significantly more than the amount they received (4.11 ± 3.58, Wilcoxon signed-rank test: z = - 2.43, p = 0.01). This may indicate that the prosocial individuals indeed cared for benefitting group members rather than gaining immediate advantages through reciprocity.

### Aggression and solicitation

We found very few instances of aggression from helpers to receivers (6.59%) after provisioning food (Goodness of fit Chi-square test: χ^2^ = 137.16, p < 0.001). Also, active solicitation in the form of interacting with the provisioning side (Pos. 1) and reaching-out behavior (i.e. holding the food pipe) were observed (16.48%) in significantly few instances by receivers toward helpers (Goodness of fit Chi-square test: χ^2^ = 81.78, p < 0.001).

Furthermore, in 22% of the trials, individuals were already present within one arm’s distance from Pos. 1, i.e. at the receiving end when a prosocial individual pressed the handle and provided food. In all other trials, i.e. in a significantly higher number of trials (Goodness of fit Chi-square test: χ^2^ = 57.16, p < 0.001), the prosocial act of pressing the handle seemed more proactive and not specific for a particular partner, which may also explain why not every press resulted in food provisioning. This is not to say that there were no links between the provider and the individual who ultimately received the reward.

### Kinship and dyadic social tolerance

To assess what might explain the likelihood and magnitude of prosociality among different dyads, we examined the effects of kinship and dyadic social tolerance. Considering the large number of participating individuals and among individual variations in prosocial tendencies, we analyzed the likelihood and number of prosocial provisioning across all potential dyads we could create with the prosocial individuals as helpers and all participating individuals as recipients. In other words, a prosocial individual could potentially help everyone. In this analysis, kinship was found to influence the magnitude (GLMM: z = 2.990, p = 0.002; kin = 3.68 ± 7.18; non-kin = 0.32 ± 1.41; **Figure 4b**) but not the likelihood (GLMM: z = 3.534, p = 0.24, **Figure 4a**) of prosocial provisioning. Dyadic social tolerance, as measured through the string-pulling task, positively predicted both the likelihood (GLMM: z = 1.610, p = 0.01; **Figure 4a**) and magnitude of prosocial provisioning (GLMM: z = 2.258, p = 0.02; **Figure 4b**), independently from the effect of kin. We did not find any effect of age, sex and rank differences on the likelihood and magnitude of prosocial provision (**Figures 4a** and **4b**).

**Figure 4.**
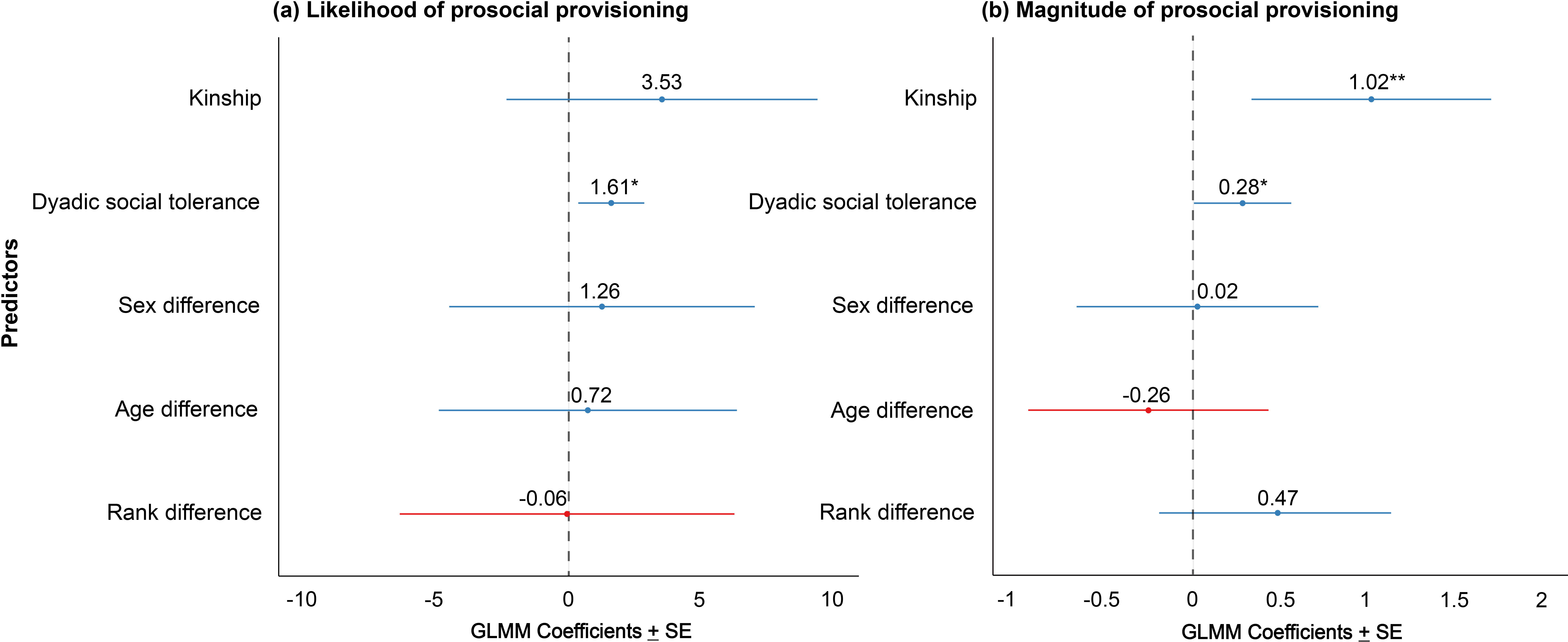
Model overview investigating the likelihood and magnitude of prosocial provisioning. GLMM coefficients ± SE of the models are presented [** *p* < 0.01, * *p* < 0.05]. Blue lines indicate positive associations between the predictors and response variable, whereas the red lines indicate negative associations; the vertical dashed line indicates no effect. (a) We found only a positive effect of dyadic social tolerance on the likelihood of prosocial provisioning (b) Both kinship and dyadic social tolerance positively predicted the magnitude of prosocial provisioning.

## Discussion

We asked whether prosociality can be favored in a despotic society and, if so, what factors necessarily regulate it. Our results show that Japanese macaques, a highly despotic species that is neither self-domesticated nor cooperatively breeding, exhibits prosociality at considerably high levels. Although we found a clear kin bias in the magnitude of prosociality, prosocial tendencies were not restricted to kin-relatives, and non-kin dyads with relatively high social tolerance also engaged in prosocial acts. Besides, the low aggression and solicitation rates in our results indicate the proactive and intentional nature of such beneficial behavior. In addition, our findings reveal substantial individual differences in prosocial tendencies.

### Genuine prosocial intent

Burkart and van Schaik (10) illustrated that to be proactively prosocial, some criteria need to be fulfilled in the group service paradigm – higher frequency of press in the test than controls, sustained or emerging provisions throughout the test condition, and provision in the absence of solicitation. The prosocial intent in our group of Japanese macaques seemed genuine as they fulfilled all the above criteria.

Evidently, in our study, individuals pressed the handle more often in the test than in the control conditions. Unlike the decreasing rates of pressing in both control conditions over the subsequent sessions, the sustained or increased rates of pressing in the test condition testify that individuals understood the contingencies of the experiment well. A faster response time to press the handle in the test than control conditions further supports the above statement. Individuals were equally motivated throughout the study, i.e. independent of test or control condition, as seen from their responses in the motivation trials (96% success). Therefore, the low press rates during control conditions were not governed by the lack of willingness to participate. Moreover, the negligible amount of aggression from actors to receivers could explain the true prosocial intent of the macaques, rather than trying to procure the food rewards themselves, albeit impossible. In addition, the prosocial individuals provided food significantly more to others than they received; thus, they were also not reliant on immediate reciprocal benefits.

The fact that active solicitation and the presence of individuals being potential receivers did not necessarily elicit prosociality also indicates its proactive nature. For instance, active solicitation, begging or signal prompting has been argued to be a stimulus enhancement mechanism of rather “reactive” prosociality (19, 31, 32). In contrast, some studies even found a negative role of signal prompting on prosociality (34–36). Given the contradicting results, the mechanisms behind these responses are not clearly understood.

### Kin favoritism and the influence of dyadic social tolerance

Kin favoritism and nepotism are integral properties of Japanese macaques and, in general, despotic societies (11, 37, 38). Our study group followed species-typical matrilineal norms, i.e., the presence of distinct hierarchical ranks and order (25, 39, 40). As hypothesized, kinship was found to influence prosociality, though it did not predict the propensity or likelihood of prosocial provisioning, suggesting a general tendency to help group members. But we did find evidence of kin relationships positively influencing the magnitude of provisioning. That is, the majority of the food provisioning was restricted to kin members. These findings align with the kin-selection theory (41), i.e., prosociality among kin members can lead to higher indirect benefits (42). Besides, even though the chances of future reciprocity (43) can be expected, the actors at the time of helping might not be certain about such returns, explaining the unselfish motives from a proximate point of view (44).

Close kin membership can be instrumental in terms of social support and interdependencies, especially in despotic societies where frequent aggression and conflicts are prevalent (40, 45–47). For instance, prosociality among kin members was also found on relatively less despotic long-tailed macaques (16). Surprisingly, our study did not find an interaction effect of kinship and dyadic social tolerance. We speculate that the dyadic social tolerance task could not fully capture the different facets of relationship quality (48), another potential factor facilitating prosociality. It could also be possible that the linear hierarchical system is at play, leading to lower social tolerance levels even among the members of the same matriline than conventionally thought.

Nevertheless, we found an effect of dyadic social tolerance in this study. Surprisingly, there is a great lack of studies with dyadic social tolerance being used as a predictor of prosociality, while it is known to have a robust predictive value in cooperation success in a range of species (31, 49–53). Below we discuss the potential sources of heightened social tolerance at dyad levels and outside kin relationships.

### The evolution of prosociality, the influence of socio-ecological conditions and the ‘interdependency hypothesis’

So far, there have been two influential hypotheses explaining the phylogenetic differences in prosociality across taxa - the self-domestication hypothesis (6) and the cooperative breeding hypothesis (3, 8, 10). Japanese macaques are neither (self-) domesticated nor cooperatively breeding. Yet, interestingly, the observed food provisioning rate (∼ 69%) of the Japanese macaques in our study resembled cooperative breeding and self-domesticated species’ prosocial provisioning (3, 5, 10), notably while using the exact same set-up and procedure. These results, together with the findings of prosociality (although used different set-up and/or procedures) among other non-domesticated species and species that lack cooperative breeding or allomaternal care (16, 18, 19, 54–56), call for additional hypotheses explaining the evolution of prosocial preferences.

Both cooperative breeding and self-domestication hypothesis, to a reasonable extent, highlight dependencies among individuals and subsequently predict high social tolerance at the individual as well as group levels. Our Japanese macaque group showed, as expected, given their despotic social structure (57), very low group-level social tolerance, but we did identify ample interdependencies among its members that may explain their prosocial tendencies. We, therefore, call for a more general ‘interdependency hypothesis’ to explain the evolutionary origins of prosocial tendencies and the phylogenetic differences observed. Note that interdependency has also been postulated as an explanatory variable for variation in inequity responses across and within species (58).

Our results also question the premises of the covariation framework for macaques or primates in general (11, 24), as it expects little to no cooperative behaviors in despotic species with low group-level tolerance. We argue here that instead, in such despotic societies, cooperation may be paramount to attain and maintain rank positions and that it, as such, creates interdependencies favoring prosocial tendencies (cf. “Machiavellian Intelligence”, (14, 15)).

### Between-group variation in Japanese macaques’ prosocial tendencies

Our results are in stark contrast with previous reports on prosociality in captive Japanese macaque (∼ 0%, (10)). Consistent with the prediction that independently breeding free or semi free-ranging primates have lower social tolerance than captive populations (59–62), our study group showed lower group-level social tolerance (0.13) than the previously tested population residing in a more restricted captive setting (0.32) (10). It must be noted that our group level tolerance measure was limited to focal individuals and not the entire population. Nevertheless, this again suggests that group-level social tolerance may not be a good predictor of prosociality. Yet, striking variation at the dyadic social tolerance level was observed among kin and non-kin group members, and as discussed above, they did predict the likelihood and magnitude of prosociality in our population. The contrasting results between the two Japanese macaque populations could potentially be attributed to the more complex socio-ecological conditions of the large semi free-ranging group, leading to variations in individual behavioral responses.

Relatively large size and complexity of a group can drive cognitive processes, as proposed by the social intelligence hypothesis (15, 63, 64), even within single species (65). Such complexity might create opportunities for individuals to become “Machiavellian” and make them take “political” decisions (63, 66). Since despotic societies are governed by support, especially during conflicts and coalitions (66, 67), high dyadic tolerance among both kin and non-kin members can emerge as a by-product. A classic example in the current study would be the beta male providing food to the beta female (**Movie S3**). Thus, maintaining alliances with other high-ranked individuals could help the beta male rank up in the hierarchy and become the alpha. Such interdependencies may favor a more flexible use of prosocial behaviour.

Finally, prosociality has been argued to be the foundation for inter-group conflict (68); consequently, in-group prosociality and parochial cooperation can be favored. Our study group recently underwent fission, forming two distinct subgroups (Hammer and Stribos et al., *unpublished data*), and all participating individuals in the current study belonged to one of the subgroups. Therefore, it would be interesting to examine in future studies whether such an ecological process of group fission may have governed the prosocial displays we witnessed in our study.

## Conclusion

We provide the first experimental report of proactive prosociality in semi free-ranging Japanese macaques. We discussed how kin favoritism and dyadic social tolerance in a despotic society could drive prosocial motivations. Although we did not find the age difference of the individuals to influence prosociality significantly, participation of a relatively large number of juveniles can not be neglected; for instance, due to their higher social tolerance, bonobo sub-adults have been shown to be more prosocial than adults (32). Nonetheless, prosociality was not restricted to juveniles, and therefore, we speculate on the role of complex socio-ecological and living conditions in facilitating individual prosocial motivations. While our findings contradict the predictions of the cooperative breeding hypothesis (also see (69, 70)), we emphasize inter-individual dependencies other than cooperative breeding to be valued and included in an overarching ‘interdependency hypothesis’, which can be systematically tested using an evolutionary framework. Finally, as previously stated by other researchers, we would also like to encourage exploring multiple groups of a species with similar and different living conditions (i.e., captive, semi free-ranging, and free-ranging.) and structures, to come to a conclusion on that species’ prosocial tendencies. Similarly, it might be interesting to see whether population differences in prosociality can be due to differential cultures with regard to societal norms in these populations (71).

## Materials and Methods

***Ethical note:*** The study was non-invasive and complied with Austrian Law (§ 2. Federal Law Gazette number 501/1989) and the Code for Best Practices in Field Primatology and received oversight from and was authorized by the internal board of the Austrian Research Center for Primatology. No invasive research or experimental procedures requiring ethics approval according to the European Directive 2010/63 were performed. Our study adhered to the American Society of Primatologists’ principles for the ethical treatment of primates and all applicable international, national and institutional guidelines for the care and use of animals.

### Group service paradigm

We adopted a modified version (5) of the group-service paradigm (10). The procedure included several phases occurring in the following order – Phase 0 (habituation to apparatus), Phase 1 (initial training and habituation to procedure), Phase 2 (food distribution assessment), Phase 3 (apparatus training), Phase 4 (group-service), Phase 5 (blocked control), and Phase 6 (Re-test). Phases 0, 1 & 3 consisted of habituation and training sessions (see **Supplemental Information** for details). It is important to note, however, that 25 individuals (adult female = 13, adult male = 3, juvenile female = 6, juvenile male = 3, **Table S2**) successfully passed the criteria of phases 1 and 3 and that the number of trials in all other phases was based on this number of participants, i.e. (N * 5 = 25 * 5 = 125 trials).

To measure social tolerance at the group level, we ran a food distribution assessment in phase 2. The seesaw mechanism was locked, and the board was tilted towards the macaques. Therefore, food rolled down to them when placed in Pos. 1. A trial ended with an individual obtaining the food; otherwise, it lasted a maximum of two minutes. We noted the identities of the individuals who obtained rewards over two sessions.

In the *test sessions* of phase 4, we assessed the prosocial tendency of the individuals. A food reward (shelled peanut) in Pos. 1 of the seesaw apparatus (see **Figure 1**) was placed while the wooden board was still titled toward the experimenter. As the reward was placed out of reach of the individuals, someone had to press the handle to operate the seesaw for the food to roll down and become available. Note that the actor could not reach the available food itself after pressing the handle since it would have to let go of the handle (Pos. 0) to reach the receiving end of Pos.1. Yet, by letting go of the handle, the seesaw mechanisms would tilt back to the original position, and the food would roll back out of reach. Thus, an individual could only make food available for their group mates. In addition to the test sessions, we conducted *empty control* sessions, identical to the test sessions, except that no food was placed, even though the same movement (of placing food in Pos. 1) was made to control for stimulus enhancement. We conducted five such *test*s and five *empty control* sessions, alternately, each containing 125 trials.

To investigate whether pressing in the test was only due to the presence of food, we also performed a *blocked control* in Phase 5. Here, the access to the potentially available food in Pos. 1 was blocked using plexiglass. As in Phase 4, we performed five blocked control sessions, alternating them with *empty (blocked) control* sessions.

Finally, in Phase 6 (Re-test), two additional *tests* and two *empty control* sessions were conducted alternately to eliminate the potential bias of the order of the phases in the experiment.

For phases 4–6, in each session, be it *test* or *control*, we added motivation trials where food was placed in Pos. 0 at regular intervals, i.e. after every fifth trial. These trials were included to ensure low pressing rates in either test or control sessions (if observed) were not due to a lack of motivation to participate. Overall, the monkeys were very motivated as they pressed in 96% of these trials, independent of the experimental conditions (Test – Empty control, Goodness of fit Chi-square test: χ^2^ = 0, p = 1; Test – Blocked control, χ^2^ = 0.709, p = 0.67; Empty control – Blocked control, χ^2^ = 0.709, p = 0.67). Thus, motivation alone did not influence pressing the handle in either condition.

All sessions were video-recorded, and the identities of the individuals (pressing, providing and receiving) and latencies to press were coded. To allow individuals to learn the contingencies of the different control conditions, we analyzed their performances in only the 4^th^ and 5^th^ sessions of each condition. See additional methods in the **Supplementary information** for details.

### String-pulling (tolerance) task

For the string-pulling (dyadic tolerance) task, we used a movable wooden platform (length ∼ 1.5 m) to which two ropes were attached at the two extremes (**Movie S2**). Two pieces of highly preferred food rewards (peanuts) were placed out of reach of the macaques on two ends of the platform (∼ 1m apart). The platform could only be moved by pulling one of the ropes. As this task did not require joint action, one individual could pull, move the platform and monopolize the two food rewards. We recorded the individuals who tolerated each other in front of such highly rewarding food items, i.e., simultaneously obtained rewards while sitting next to each other during a trial. A total of 18 sessions were conducted, each consisting of 20 trials.

See additional methods in the **Supplementary Information** section for detailed descriptions of study subjects, experimental procedure, coding and data analyses.

## Acknowledgements and funding sources

We are thankful to the Affenberg Zoobetriebsgesellschaft mbH, and specifically to its directors Peter and Svenja Gaubatz, for allowing us to do our research there. We also thank all animal caretakers for their excellent care of our study animals and especially Max Dorner for his help with the building of the apparatuses. The project has received funding from the European Union’s Horizon 2020 research and innovation programme under grant number H2020-MSCA-IF-2019-893016. The funder had no role in study design, data collection and analysis, publication decision, or manuscript preparation.

## Reference

1. N. Eisenberg, R. A. Fabes, T. L. Spinrad, “Prosocial Development” in Handbook of Child Psychology, (John Wiley & Sons, Inc., 2007) https:/doi.org/10.1002/9780470147658.chpsy0311.

2. J. B. Silk, Empathy, sympathy, and prosocial preferences in primates (Oxford University Press, 2007) https:/doi.org/10.1093/oxfordhb/9780198568308.013.0010.

3. J. M. Burkart, et al., The evolutionary origin of human hyper-cooperation. Nature Communications 5, 4747 (2014).

4. K. A. Cronin, Prosocial behaviour in animals: the influence of social relationships, communication and rewards. Animal Behaviour 84, 1085–1093 (2012).

5. L. Horn, et al., Sex-specific effects of cooperative breeding and colonial nesting on prosociality in corvids. Elife 9 (2020).

6. B. Hare, V. Wobber, R. Wrangham, The self-domestication hypothesis: evolution of bonobo psychology is due to selection against aggression. Animal Behaviour 83, 573– 585 (2012).

7. F. E. de Oliveira Terceiro, M. de F. Arruda, C. P. van Schaik, A. Araújo, J. M. Burkart, Higher social tolerance in wild versus captive common marmosets: the role of interdependence. Scientific Reports 11, 825 (2021).

8. J. M. Burkart, S. B. Hrdy, C. P. van Schaik, Cooperative breeding and human cognitive evolution. Evolutionary Anthropology: Issues, News, and Reviews 18, 175– 186 (2009).

9. L. Horn, C. Scheer, T. Bugnyar, J. J. M. Massen, Proactive prosociality in a cooperatively breeding corvid, the azure-winged magpie (*Cyanopica cyana*). Biology Letters 12, 20160649 (2016).

10. J. M. Burkart, C. van Schaik, Group service in macaques (Macaca fuscata), capuchins (Cebus apella) and marmosets (Callithrix jacchus): A comparative approach to identifying proactive prosocial motivations. Journal of Comparative Psychology 127, 212–225 (2013).

11. B. Thierry, Where do we stand with the covariation framework in primate societies? American Journal of Biological Anthropology 178, 5–25 (2022).

12. B. Thierry, Unity in diversity: Lessons from macaque societies. Evolutionary Anthropology: Issues, News, and Reviews 16, 224–238 (2007).

13. G. Schino, Grooming and agonistic support: a meta-analysis of primate reciprocal altruism. Behavioral Ecology 18, 115–120 (2007).

14. D. Maestripieri, Macachiavellian Intelligence (University of Chicago Press, 2007) https:/doi.org/10.7208/chicago/9780226501215.001.0001.

15. A. Whiten, R. W. Byrne, “The Machiavellian Intelligence Hypothesis” in Machiavellian Intelligence: Social Expertise and the Evolution of Intellect in Monkeys, Apes, and Humans, (Clarendon Press/Oxford University Press, 1988), pp. 1–9.

16. J. J. M. Massen, L. M. van den Berg, B. M. Spruijt, E. H. M. Sterck, Generous Leaders and Selfish Underdogs: Pro-Sociality in Despotic Macaques. PLoS ONE 5, e9734 (2010).

17. S. F. Brosnan, et al., Competing demands of prosociality and equity in monkeys. Evolution and Human Behavior 31, 279–288 (2010).

18. E. J. C. van Leeuwen, et al., Chimpanzees behave prosocially in a group-specific manner. Science Advances 7 (2021).

19. A. P. Melis, et al., Chimpanzees help conspecifics obtain food and non-food items. Proceedings of the Royal Society B: Biological Sciences 278, 1405–1413 (2011).

20. F. Warneken, M. Tomasello, Altruistic Helping in Human Infants and Young Chimpanzees. Science (1979) 311, 1301–1303 (2006).

21. C. Tennie, K. Jensen, J. Call, The nature of prosociality in chimpanzees. Nature Communications 7, 13915 (2016).

22. J. B. Silk, et al., Chimpanzees are indifferent to the welfare of unrelated group members. Nature 437, 1357–1359 (2005).

23. F. Amici, E. Visalberghi, J. Call, Lack of prosociality in great apes, capuchin monkeys and spider monkeys: convergent evidence from two different food distribution tasks. Proceedings of the Royal Society B: Biological Sciences 281, 20141699 (2014).

24. B. Thierry, Covariation of Conflict Management Patterns across Macaque Species. Natural Conflict Resolution (2000) https:/doi.org/10.1006/anbe.2001.1656.

25. B. Chapais, C.-E. G. St-Pierre, Kinship Bonds Are Not Necessary for Maintaining Matrilineal Rank in Captive Japanese Macaques. International Journal of Primatology 18, 375–385 (1997).

26. R. Ventura, B. Majolo, N. F. Koyama, S. Hardie, G. Schino, Reciprocation and interchange in wild Japanese macaques: grooming, cofeeding, and agonistic support. American Journal of Primatology 68, 1138–1149 (2006).

27. T. Kawazoe, S. Sosa, Social networks predict immigration success in wild Japanese macaques. Primates 60 (2019).

28. B. Chapais, J. Prud’Homme, S. Teijeiro, Dominance competition among siblings in Japanese macaques: Constraints on nepotism. Animal Behaviour 48 (1994).

29. Y. Kaigaishi, M. Nakamichi, K. Yamada, High but not low tolerance populations of Japanese macaques solve a novel cooperative task. Primates 60, 421–430 (2019).

30. R. Sigmundson, et al., Exploring the Cognitive Capacities of Japanese Macaques in a Cooperation Game. Animals 11, 1497 (2021).

31. J. S. Martin, S. E. Koski, T. Bugnyar, A. V. Jaeggi, J. J. M. Massen, Prosociality, social tolerance and partner choice facilitate mutually beneficial cooperation in common marmosets, Callithrix jacchus. Animal Behaviour 173, 115–136 (2021).

32. J. Verspeek, E. J. C. van Leeuwen, D. W. Laméris, N. Staes, J. M. G. Stevens, Adult bonobos show no prosociality in both prosocial choice task and group service paradigm. PeerJ 10, e12849 (2022).

33. E. C. Pielou, The measurement of diversity in different types of biological collections. Journal of Theoretical Biology 13, 131–144 (1966).

34. A. Takimoto, H. Kuroshima, K. Fujita, Capuchin monkeys (Cebus apella) are sensitive to others’ reward: an experimental analysis of food-choice for conspecifics. Animal Cognition 13, 249–261 (2010).

35. K. A. Cronin, K. K. E. Schroeder, E. S. Rothwell, J. B. Silk, C. T. Snowdon, Cooperatively breeding cottontop tamarins (Saguinus oedipus) do not donate rewards to their long-term mates. Journal of Comparative Psychology 123, 231–241 (2009).

36. J. M. Burkart, A. Heschl, Understanding visual access in common marmosets, Callithrix jacchus: perspective taking or behaviour reading? Animal Behaviour 73, 457–469 (2007).

37. J. B. Silk, Nepotistic cooperation in non-human primate groups. Philosophical Transactions of the Royal Society B: Biological Sciences 364, 3243–3254 (2009).

38. T. Matsuzawa, W. C. McGrew, Kinji Imanishi and 60 years of Japanese primatology. Current Biology 18, R587–R591 (2008).

39. L. S. Pflüger, et al., Twenty-three-year demographic history of the Affenberg Japanese macaques (Macaca fuscata), a translocated semi-free-ranging group in southern Austria. Primates 62, 761–776 (2021).

40. B. Chapais, Rank Maintenance in Female Japanese Macaques: Experimental Evidence for Social Dependency. Behaviour 104, 41–58 (1988).

41. W. D. Hamilton, The genetical evolution of social behaviour. II. Journal of Theoretical Biology 7, 17–52 (1964).

42. F. B. M. de Waal, K. Leimgruber, A. R. Greenberg, Giving is self-rewarding for monkeys. Proceedings of the National Academy of Sciences 105, 13685–13689 (2008).

43. R. L. Trivers, The Evolution of Reciprocal Altruism. The Quarterly Review of Biology 46, 35–57 (1971).

44. F. B. M. de Waal, M. Suchak, Prosocial primates: selfish and unselfish motivations. Philosophical Transactions of the Royal Society B: Biological Sciences 365, 2711– 2722 (2010).

45. I. Norscia, E. Palagi, The socio-matrix reloaded: from hierarchy to dominance profile in wild lemurs. PeerJ 3, e729 (2015).

46. E. Palagi, I. Norscia, The Season for Peace: Reconciliation in a Despotic Species (Lemur catta). PLOS ONE 10, e0142150 (2015).

47. K. Watanabe, Alliance formation in a free-ranging troop of Japanese macaques. Primates 20, 459–474 (1979).

48. M. Cords, Filippo Aureli, “Relationship Qualities” in Natural Conflict Resolution, F. B. M. de Waal, Ed. (University of California Press, 2000), pp. 177–198.

49. S. Molesti, B. Majolo, Cooperation in wild Barbary macaques: factors affecting free partner choice. Animal Cognition 19, 133–146 (2016).

50. J. J. M. Massen, C. Ritter, T. Bugnyar, Tolerance and reward equity predict cooperation in ravens (Corvus corax). Scientific Reports 5, 15021 (2015).

51. B. Hare, A. P. Melis, V. Woods, S. Hastings, R. Wrangham, Tolerance Allows Bonobos to Outperform Chimpanzees on a Cooperative Task. Current Biology 17, 619–623 (2007).

52. A. P. Melis, B. Hare, M. Tomasello, Engineering cooperation in chimpanzees: tolerance constraints on cooperation. Animal Behaviour 72, 275–286 (2006).

53. D. Werdenich, L. Huber, Social factors determine cooperation in marmosets. Animal Behaviour 64, 771–781 (2002).

54. M. J. M. Gachomba, et al., Multimodal cues displayed by submissive rats promote prosocial choices by dominants. Current Biology (2022) https:/doi.org/10.1016/j.cub.2022.06.026.

55. I. Ben-Ami Bartal, et al., Neural correlates of ingroup bias for prosociality in rats. Elife 10 (2021).

56. J. J. M. Massen, I. J. A. F. Luyten, B. M. Spruijt, E. H. M. Sterck, Benefiting friends or dominants: prosocial choices mainly depend on rank position in long-tailed macaques (Macaca fascicularis). Primates 52, 237–247 (2011).

57. P. M. Kappeler, A framework for studying social complexity. Behavioral Ecology and Sociobiology 73, 13 (2019).

58. S. F. Brosnan, A hypothesis of the co-evolution of cooperation and responses to inequity. Frontiers in Neuroscience (2011) https:/doi.org/10.3389/fnins.2011.00043.

59. F. Amici, A. Widdig, L. von Fersen, A. Lopez Caicoya, B. Majolo, Intra-specific Variation in the Social Behavior of Barbary macaques (Macaca sylvanus). Frontiers in Psychology 12 (2021).

60. C. Fichtel, A. v. Schnoell, P. M. Kappeler, Measuring social tolerance: An experimental approach in two lemurid primates. Ethology 124, 65–73 (2018).

61. C. P. van Schaik, C. H. Janson, Recognizing the Many Faces of Primate Food Competition: Methods. Behaviour 105, 165–186 (1988).

62. L. A. Isbell, Contest and scramble competition: patterns of female aggression and ranging behavior among primates. Behavioral Ecology 2, 143–155 (1991).

63. R. I. M. Dunbar, The social brain hypothesis. Evolutionary Anthropology: Issues, News, and Reviews 6, 178–190 (1998).

64. N. K. Humphrey, “The social function of intellect” in Growing Points in Ethology, (1976), pp. 303–317.

65. B. J. Ashton, A. R. Ridley, E. K. Edwards, A. Thornton, Cognitive performance is linked to group size and affects fitness in Australian magpies. Nature 554 (2018).

66. F. B. M. de Waal, L. M. Luttrell, Mechanisms of social reciprocity in three primate species: Symmetrical relationship characteristics or cognition? Ethology and Sociobiology 9, 101–118 (1988).

67. C. K. Hemelrijk, I. Puga-Gonzalez, An Individual-Oriented Model on the Emergence of Support in Fights, Its Reciprocation and Exchange. PLoS ONE 7, e37271 (2012).

68. C. K. W. de Dreu, A. Fariña, J. Gross, A. Romano, Prosociality as a foundation for intergroup conflict. Current Opinion in Psychology 44 (2022).

69. A. Thornton, K. McAuliffe, Cognitive consequences of cooperative breeding? A critical appraisal. Journal of Zoology 295, 12–22 (2015).

70. A. Thornton, et al., Fundamental problems with the cooperative breeding hypothesis. A reply to Burkart & van Schaik. Journal of Zoology 299, 84–88 (2016).

71. B. R. House, et al., Universal norm psychology leads to societal diversity in prosocial behaviour and development. Nature Human Behaviour 4 (2020).

