## Supplementary Information for "Prosociality in a despotic society"

----------------------------------------------------------------------------------------------------------------

* Correspondance:

**Table of contents:**

**Page**

**Table S1:** List of participating individuals. 2

**Figure S1.** Food distribution during phase 2 of the group service paradigm. 3

**Table S2.** Overview of individuals pressing the handle. 4

**Figure S2.** Latency of pressing the handle in different conditions. 5

**Figure S3.** Overview of food provision by individuals. 6

**Table S3.** Number of food provided and received in the test condition. 7

**Additional methods**	8-13

**Media file descriptions**	13

**References**	14

**Table S1. List of participating individuals.** Individuals who participated in the study voluntarily, out of the ~170 individuals present in the population.

| **Individuals** | **Age (yo)** | **Age group** | **Year of birth** | **Sex** | **Matriline rank** |
| --- | --- | --- | --- | --- | --- |
| Andy | 1 | Juvenile | 2021 | ♂ | 10 |
| Cinderella | 2 | Juvenile | 2022 | ♀ | 7 |
| Clara | 5 | Adult | 2017 | ♀ | 10 |
| Dornroschen | 2 | Juvenile | 2020 | ♀ | 5 |
| Frau Holle | 2 | Juvenile | 2020 | ♀ | 4 |
| Frida | 1 | Juvenile | 2021 | ♀ | 1 |
| Fuji | 3 | Juvenile | 2019 | ♂ | 5 |
| Goldmarie | 2 | Juvenile | 2020 | ♀ | 7 |
| Herta | 10 | Adult | 2012 | ♀ | 5 |
| Iris **(α)** | 21 | Adult | 2001 | ♀ | 1 |
| Janis | 8 | Adult | 2014 | ♀ | 7 |
| Jessy **(β)** | 12 | Adult | 2010 | ♀ | 1 |
| Kate | 11 | Adult | 2011 | ♀ | 1 |
| Kiki | 1 | Juvenile | 2021 | ♀ | 2 |
| Krato | 9 | Adult | 2013 | ♀ | 5 |
| Lisa | 11 | Adult | 2001 | ♀ | 5 |
| Marie | 5 | Adult | 2017 | ♀ | 4 |
| Pauli | 22 | Adult | 2000 | ♂ | 2 |
| Pippi | 6 | Adult | 2016 | ♀ | 2 |
| Salvador | 1 | Juvenile | 2021 | ♂ | 4 |
| Sandra | 11 | Adult | 2011 | ♀ | 4 |
| Spooky **(α)** | 10 | Adult | 2012 | ♂ | 3 |
| Uschi | 10 | Adult | 2012 | ♀ | 7 |
| Wicky **(β)** | 10 | Adult | 2012 | ♂ | 6 |
| Zarah | 7 | Adult | 2015 | ♀ | 7 |

*The matriline number refers to the matriline rank and dominance already assessed for this group* (1)*. All individuals from the same matriline are related to a certain degree and thus are kin-relatives. The alpha and beta (males and females) are indicated. Individuals older than four years old were considered adults, and individuals younger than four were juveniles* (2)*.*

**Figure S1. Food distribution during phase 2 of the group service paradigm.**

No. of food rewards obtained (Social tolerance phase)

| Individuals | Food |
| --- | --- |
| Wicky | 71 |
| Spooky | 46 |
| Iris | 29 |
| Sandra | 12 |
| Pippi | 11 |
| Dornroschen | 2 |
| Jessy | 2 |
| Kiki | 2 |
| Frida | 1 |
| Goldmarie | 1 |
| Andy | 0 |
| Cinderella | 0 |
| Clara | 0 |
| Frau Holle | 0 |
| Fuji | 0 |
| Herta | 0 |
| Janis | 0 |
| Kate | 0 |
| Krato | 0 |
| Marie | 0 |
| Pauli | 0 |
| Salvador | 0 |
| Uschi | 0 |
| Zarah | 0 |
| Lisa | 0 |

*A total of 250 food rewards were placed over two sessions, out of which 177 rewards were obtained by the focal individuals (non-focal individuals were not included in the analysis, i.e., only the 25 participating monkeys were considered). See R-script for the procedure to calculate Pielou's J′ (obtained value = 0.13, suggesting low group-level social tolerance).*

**Table S2. Overview of individuals pressing the handle in sessions 4 and 5 of the group service paradigm.**

|  | Sessions 4 & 5 | | |
| --- | --- | --- | --- |
| **Individuals** | **Test** | **Empty control** | **Blocked control** |
| Andy | 1 | 0 | 7 |
| *Cinderella* | 15 | 23 | 0*** |
| Clara | 1 | 3 | 3 |
| *Dornroschen* | 85 | 4*** | 3*** |
| *Frau Holle* | 9 | 0** | 5 |
| *Frida* | 22 | 3*** | 13 |
| *Fuji* | 17 | 0*** | 1*** |
| Goldmarie | 2 | 5 | 0 |
| Herta | 5 | 4 | 5 |
| Iris | 0 | 2 | 0 |
| Janis | 1 | 7 | 4 |
| *Jessy* | 10 | 0** | 3 |
| Kate | 8 | 3 | 5 |
| Kiki | 4 | 0 | 5 |
| Krato | 4 | 5 | 6 |
| Lisa | 3 | 0 | 2 |
| *Marie* | 19 | 20 | 4** |
| Pauli | 4 | 0 | 2 |
| Pippi | 0 | 3 | 0 |
| Salvador | 1 | 3 | 5 |
| *Sandra* | 9 | 0** | 8 |
| Spooky | 6 | 1 | 2 |
| Uschi | 0 | 0 | 0 |
| *Wicky* | 20 | 5** | 2*** |
| *Zarah* | 7 | 0* | 2 |

**Fisher’s exact tests**: *** *p* < 0.001, ** *p* < 0.01, * *p* < 0.05.

*Names in italics represent prosocial individuals. Note that all individuals passed the criterion of phases 1 and 3. We compared test-empty control and test-blocked control presses for every individual using Fisher’s exact tests with Bonferroni correction.*

**Figure S2**

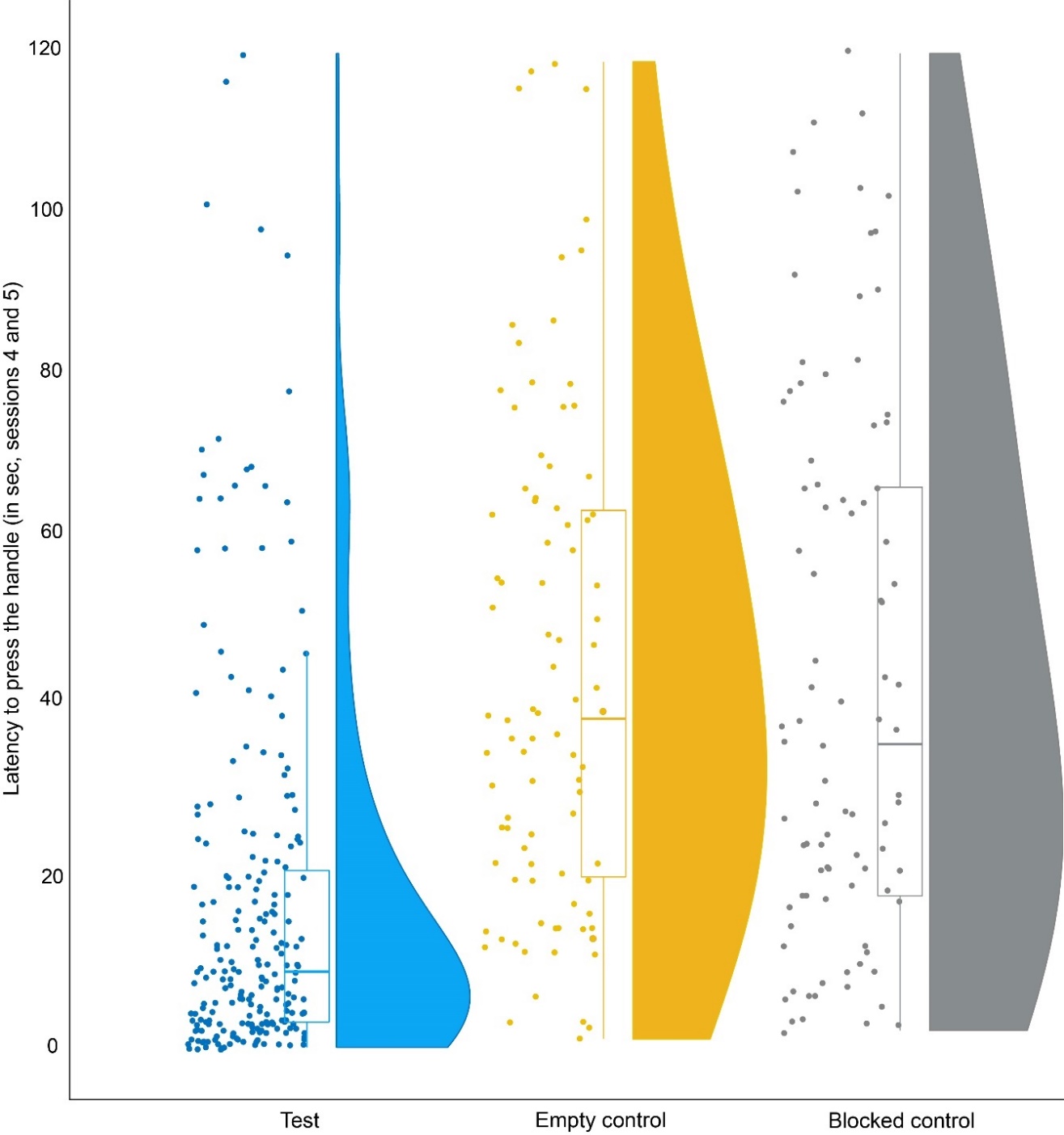

**Figure S2. Latency of pressing the handle in different conditions.** *Individuals were faster at pressing the handle in the test (18.32 ± 22.31 sec) than in empty (45.37 ± 29.18 sec, LM: t = 9.584, p < 0.001) and blocked (44.52 ± 32.18 sec, LM: t = 9.047, p < 0.001) control conditions. Half-violin plots indicate the distribution, while solid dots indicate the raw values. The boxes illustrate the interquartile range, horizontal bars inside the boxes indicate median values and whiskers indicate the range of the data.*

**Figure S3**

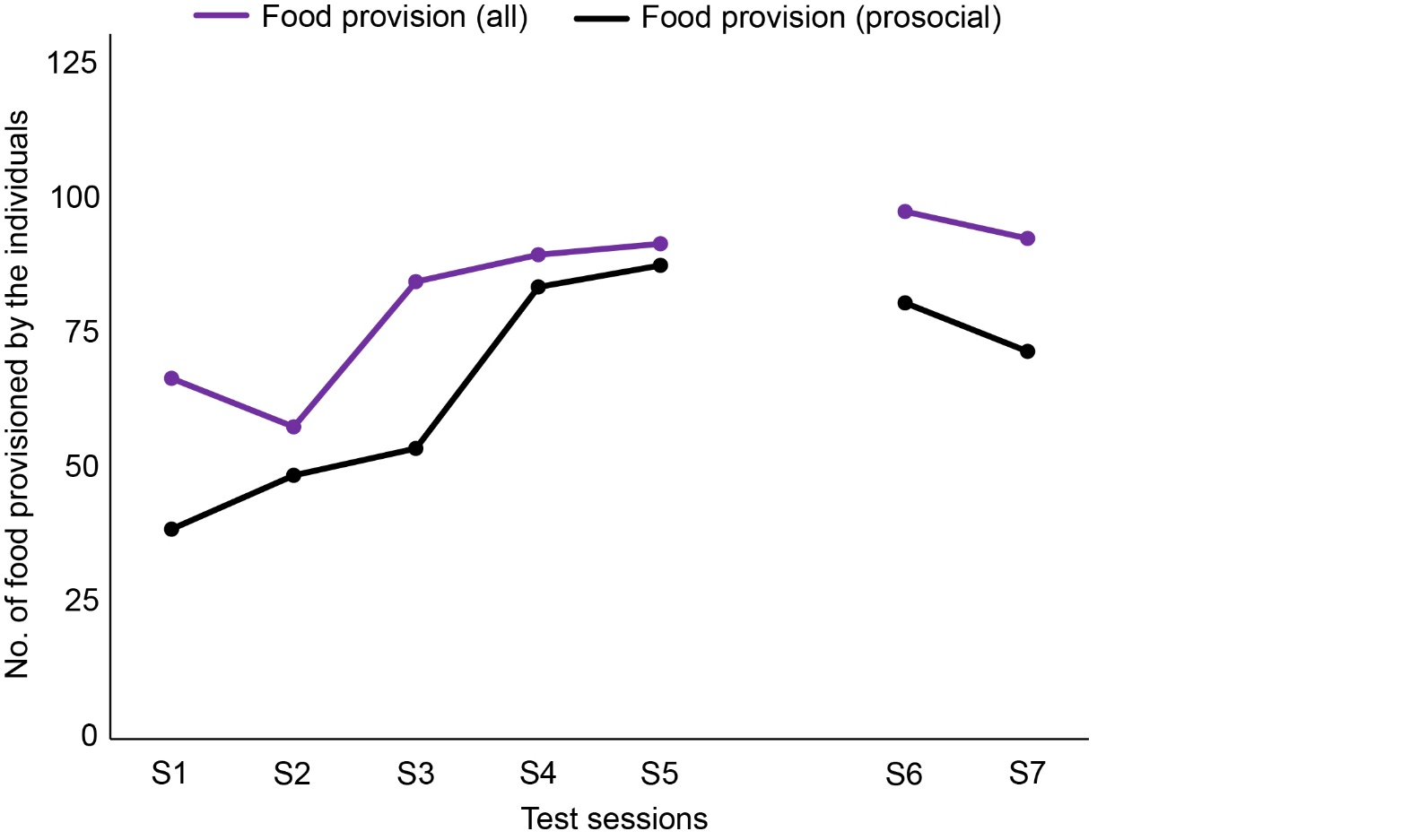

**Figure S3. Overview of food provision by individuals across all sessions in the test condition.** *S1-S5: regular sessions, while S6 and S7 indicate re-test sessions. The prosocial individuals provisioned food at a sustained rate (S1: 31%; S2: 39%; S3: 43%; S4: 67%; S5: 70%). A significant difference, however, was noticed between session one (no. of provisions ± SD: 3.9 ± 9.02) and five (8.8 ± 9.67) of the test condition (GLMM: z = 2.478, p = 0.013), indicating an increased food provision from session one to five and suggesting better coordination between actors and recipients. Food provisions in the re-test sessions were comparable (S6: 65%; S7: 58%) to earlier sessions, indicating no bias due to the pre-determined order of the experimental phases.*

**Table S3. Number of food provided and received during S4 and S5 phases of the test condition of the group service paradigm.**

| Individuals | Provided | Received |
| --- | --- | --- |
| Andy | 0 | 0 |
| *Cinderella* | 11 | 3 |
| Clara | 1 | 3 |
| *Dornroschen* | 78 | 9 |
| *Frau Holle* | 9 | 7 |
| *Frida* | 22 | 1 |
| *Fuji* | 17 | 5 |
| Goldmarie | 1 | 3 |
| Herta | 0 | 1 |
| Iris | 0 | 0 |
| Janis | 1 | 5 |
| Jessy | 0 | 28 |
| Kate | 2 | 6 |
| Kiki | 2 | 25 |
| Krato | 2 | 10 |
| Lisa | 0 | 41 |
| *Marie* | 15 | 3 |
| Pauli | 1 | 2 |
| Pippi | 0 | 0 |
| Salvador | 0 | 10 |
| *Sandra* | 7 | 9 |
| Spooky | 0 | 0 |
| Uschi | 0 | 0 |
| *Wicky* | 11 | 0 |
| *Zarah* | 2 | 0 |

*We compared the number of food provisions and food received for the prosocial and non-prosocial individuals separately. Prosocial individuals provided food to others (19.11 ± 22.83) significantly more than the amount of food they received (4.11 ± 3.58, Wilcoxon signed-rank test: z = - 2.43, p = 0.01). Note that one individual (Jessy), although pressed more in the test than in empty control, never provided food to others. We, therefore, excluded her from the list of prosocial individuals for a conservative measure.*

**Additional methods:**

**(a) Study site and subjects:** The study was conducted with a semi-free-ranging group (~170 individuals) of Japanese macaques at Affenberg Landskron in Austria from February to May 2022. The macaques live in natural socio-environmental conditions with limited human intervention, except for feeding and emergency medical purposes (such as veterinary care). However, the group is habituated to human presence as visitors can enter the enclosure within the scope of guided tours offered by the park from April to September. Our study gained high ecological validity due to the population's living conditions and naturalistic group setting (**Figure S4**).

**Figure S4 -**

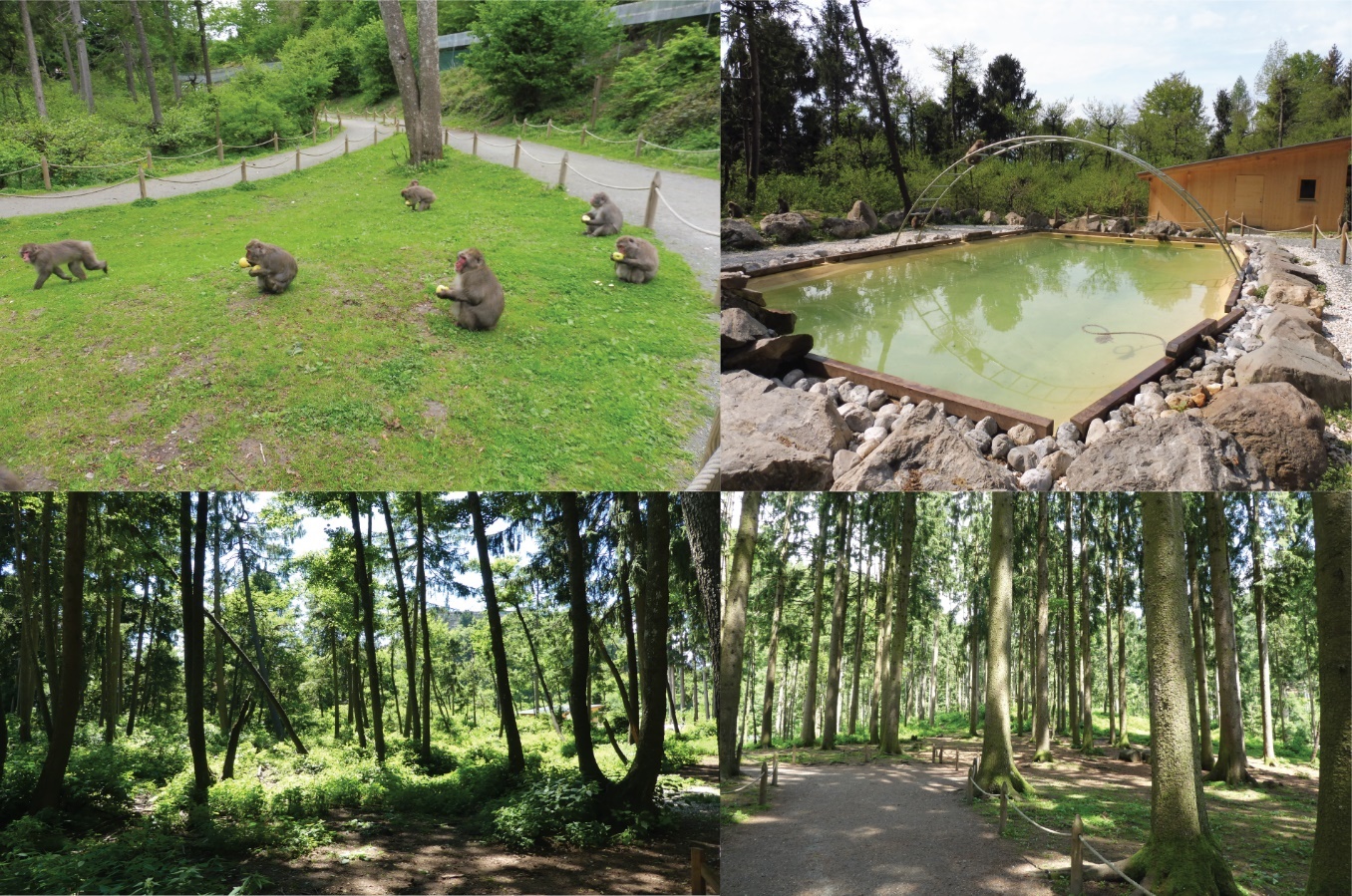

**Figure S4. Different sections of the Affenberg Landskron.** *Photos by Roy Hammer.*

A total of 25 self-trained individuals voluntarily participated in the study while remaining in their social group. Direct interaction between the subjects and the experimenter was avoided by placing the apparatuses needed for our experiments in small research huts (with wire mesh separations), which the macaques could operate from outside. Affenberg Landskron provided age, sex, and rank data (see raw data sheet “large”) of the individuals as long-term behavioural observations were being conducted for other studies with the population (see **Table S1**).

**(b) Apparatuses:** We used a seesaw mechanism for the group service paradigm (3, 4) , whereas a string-pulling task was conducted to measure dyadic social tolerance (5). The seesaw tool used a wooden board (length ~ 1.5 m) attached to the grids of the wire mesh separation of a research hut. The wooden board had two transparent plastic pipes (Ø ~ 3 inches) attached to two extremes (pos 0 and pos 1) through which food rewards (i.e., peanuts) could move. By default, the board was tilted towards the experimenter. A metal handle was connected to one of the ends of the board (pos 0) and projected outside, which upon pressing, could tilt the board towards the macaques, resulting in food rolling down through the pipe(s) (if present). In an attempt to press or when an individual released the handle halfway, the seesaw moved back to its original position, and the food (if present) would role out of reach again. Therefore, it was essential to press fully and hold the handle to get the food rewards.

For the string-pulling (dyadic tolerance) task, we used a wooden platform (length ~ 1.5 m) to which two strings were attached at the two extremes. Food rewards were placed out of reach of the macaques on two ends of the platform, which could only be moved by pulling one of the strings. The task required no joint action. The length of the apparatuses was kept constant.

**(c) Experimental Procedure:**

**(i) Group service paradigm –** The protocol consisted of several phases: habituation to the apparatus, social tolerance phase, apparatus training, group service (test and empty control), and blocked controls in a pre-determined order (see below). Each phase served a purpose and came with a criterion that needed to be fulfilled to begin the next phase. The rewards used were peanuts.

***Phase 0 (Habituation phase)***

This phase was carried out to reduce and/or eliminate any potential neophobia towards the seesaw apparatus and to habituate individuals to the setup. Food items were spread over the wooden board at regular intervals. The experimenter tried to catch the attention of the individuals by calling “Monkeys”. Three sessions were conducted (each on one day for ~1 hour). A total of 25 individuals participated in this phase voluntarily and obtained food rewards successfully.

***Phase 1 (Initial training and habituation to procedure)***

The mechanism of the seesaw was completely locked, and the platform was tilted towards the individuals. So, when placed in either position, food rewards would roll down to the individuals. We conducted a total of five sessions, and for each session, food was provided in pos. 0 and pos. 1, alternately. The number of trials was determined according to the number of individuals who successfully passed phase 0 (25 * 5 = 125 trials/session, see (6)). This number was used for the following phases as well. A trial began when a food reward was placed (either in Pos. 0 or Pos. 1) and ended with an individual retrieving it or after 2 min. In this phase, an individual was considered trained if they received at least a total of 10 rewards throughout the different sessions. To fulfil the criterion, at least half of the habituated individuals needed to get trained. We recorded which individual(s) obtained the rewards for each trial.

***Phase 2 (Social tolerance test)***

The procedure was the same as in phase 1, with the mechanism still locked and the platform tilted toward the individuals. However, food rewards were always placed in Pos. 1 in this phase. We conducted two sessions, with each having 125 trials.

***Phase 3 (Apparatus training)***

The seesaw mechanism was operational from this phase and onwards. Therefore, individuals needed to learn to press the handle to tilt the board and get the rewards. Food rewards were always placed in Pos. 0. Thus, individuals pressing and holding the handle could make food rewards available to themselves. The phase was completed when at least half of the individuals obtained at least ten rewards over five sessions.

***Phase 4 (group service: test and empty control)***

This phase was the core of the experiment. In *test* sessions, food was placed in Pos. 1 for each of the 125 trials of the test sessions. An individual needed to press the handle in Pos. 0 to make food accessible for another individual in Pos.1. The trial began when the experimenter placed a peanut in the pipe in Pos. 1 while drawing the attention of the macaques (by calling “Monkeys”). An actor could provision by pressing and holding the handle within a maximum duration of 2 min. If not, the experimenter removed the reward from the pipe and started a new trial. Trials were conducted at different times throughout the day, from 0930 to 1830 hours. We carried out *empty control* sessions alternately with the test sessions. Here, the experimenter pretended to put a reward in Pos. 1 but left the pipe empty to check for potential stimulus enhancement effects. The maximum duration of a trial was again 2 min.

***Phase 5 (Blocked controls)***

The access to the food pipe in Pos. 1 was blocked by attaching plexiglass to the enclosure. Food reward was thus visible but not accessible to the individuals, even if actors pressed the handle. We conducted five blocked test sessions during which food was placed in Pos. 1 (similarly to test sessions in phase 4), alternately with five empty (blocked) control sessions (no food placed in Pos. 1).

***Phase 6 (Re-test and empty controls)***

To check for any potential bias due to the experimental order (6, 7), we repeated two tests and two empty control sessions. The procedure was exactly the same as the regular test and empty control sessions.

We recorded the identities of the individuals pressing, provisioning, and receiving for all the above phases (wherever applicable). Also, we measured the latencies for pressing the handle for all the trials. For phases 4, 5 and 6, every fifth trial was a motivation trial. Food was placed in Pos. 0, accessible to the actor (if pressed). These motivation trials kept the monkeys interested in participating in the experiment. Additionally, these trials helped check if low pressing rates (if any) resulted from a general lack of willingness to participate in the experiment.

**(ii) String-pulling (dyadic tolerance) task** – The string-pulling task was designed to assess dyadic social tolerance. This task was independent of the group service paradigm and was performed afterwards.

***Habituation to apparatus***

We first conducted a habituation phase for a whole day (between 0930 and 1830 hours). We put food rewards on the platform and moved it slowly toward the monkeys. This phase aimed to familiarize the monkeys with the apparatus. Individuals were considered habituated after obtaining at least five rewards during the habituation phase.

***Dyadic social tolerance***

The experiment lasted three days, and we conducted 18 sessions of 20 trials each. Every trial began by calling “Monkeys” while placing food rewards (peanut) on either end of the platform yet out of reach of the monkeys. Once the monkeys were attentive, we presented both the strings at the same time. To access the rewards, at least one individual needed to pull. The trial ended when the platform was moved and rewards obtained or after 2 min. When the platform was moved, two pieces of food rewards were accessible, placed ~1 m apart, meaning that a high social tolerance was necessary for two individuals to retrieve the rewards simultaneously. Thus, monopolization of both rewards by a single individual was also possible.

We recorded all the phases of the two experiments using a Canon HF G86 video camera mounted on a tripod. Additionally, we noted the identities of the individuals participating using a pen and a notebook.

**(d) Data analyses**:

Identities of the individuals were noted live with regard to pressing the handle, providing and receiving food rewards by the experimenter and confirmed later via videos. A second rater, who was not the experimenter, scored the behavioral variables for all the sessions. Interrater reliability was excellent (ICC(3,k) = 0.98). For detailed step-by-step analyses and reproducibility, see **Raw data** and **R-Script**.

***Group and dyadic social tolerance***

We calculated the group-level social tolerance (refers to the focal animals and not the entire population) by calculating the evenness of reward distribution among the individuals with Pielou’s *J’* (expressed within a range between 0–1; 0 indicating maximum inequality and 1 suggesting a completely equal distribution) indicator (8).

Dyadic social tolerance was measured from the string-pulling task. The strength of dyadic social tolerance was defined by the instances where two individuals retrieved food rewards simultaneously.

***Group and individual-level press***

A Generalized linear mixed-effect model (GLMM) with Poisson distribution (log-link function) was conducted to investigate if the overall press (count data) was higher in the test than in control conditions (sessions 4 and 5). We, thus, included experimental conditions as fixed effects and individuals as random effects in the model. For individual-level press, we compared test-empty control and test-blocked control presses for every individual, using Fisher’s exact tests with Bonferroni correction.

***Sustained press and food provision across test sessions***

We used a negative binomial GLMM to check whether the individuals pressed the handle at sustaining rates (count) across the five test sessions of the group service paradigm. Another negative binomial GLMM investigated the number of food rewards provided (count) by the prosocial individuals across the five test sessions. In both models, sessions were included as categorical fixed effects, whereas individuals as random effects.

Note, since successful food provisioning (see below) depended not only on an individual’s press but also on the temporal and spatial coordination between actor and recipient, we could not exclude that a lack of coordination prevented food provisioning in some cases. Therefore, we used the sum of the number of presses in the last two sessions of the prosocial test (phase 4) of all individuals. However, for one individual (*Jessy*), no evidence of actual provisioning was observed. Thus, we excluded her from the list of prosocial individuals for a more conservative approach.

***Latency to press the handle***

We conducted a linear model (LM) analysis to investigate the effect of experimental conditions (test/empty control/blocked control) and sessions (sessions 4 and 5) on the latencies to press the apparatus handle. We considered trials with presses for this analysis.

***Effect of kinship and dyadic social tolerance on prosocial provisioning***

We looked at the effects of kinship and dyadic social tolerance on the likelihood and magnitude of prosocial provisioning. We used a hurdle model approach, a two-step process where the first model deals with zero-inflated count data, and a subsequent model examines only instances with a positive count. In addition, we included age, sex, and rank differences between the helpers (prosocial individuals) and receivers as control variables in the models. The possible combinations of dyads were built considering the identified prosocial individuals. Each of the nine identified prosocial individuals had a chance to provide food for others (possible dyads = 9 * 24 = 216; non-kin dyads = 188, kin dyads = 28).

In the first step, we used a binomial GLMM (food provision: yes/no) with kinship (yes/no) and dyadic social tolerance (count) as fixed effects. Sex, age, and rank differences (all categorical) were included as control variables. All possible combinations of dyads were built and included as random effects (i.e., actor/receiver). This model was aimed at understanding the likelihood of prosocial provisioning. The follow-up Poisson GLMM included instances where food provisions occurred (count data, i.e., food provision > 0). All other variables were the same as in the first model. We first checked for interaction effects between the two main predictors (i.e., kinship and dyadic social tolerance) for both models. In case of no significant effect, we examined the individual effects of the predictors.

All statistical analyses were carried out in R (3.6.1) (9). The GLMM and LM analyses were conducted using “lme4” package (10). Null vs full model comparisons were made for all the models “lmtest” (null vs full model comparisons, (11)). We investigated the model residual distribution, dispersion and outliers (“DHARMa”, (12)). Multicollinearity of the predictors (hurdle model, follow-up step) was examined (“car”, (13)), and a variance inflation factor (VIF) of <3 was set as a threshold for low correlation between the predictors. The significance value (α) was set as 0.05 for all statistical tests.

**Media file descriptions**

**Movie S1**

**Group-level social tolerance test during Phase 2 of the group service paradigm.** Individuals were obtaining food rewards from Pos. 1 of the apparatus. The alpha male approached and displaced others and started monopolizing.

**Movie S2**

**Dyadic social tolerance measure from the string-pulling task.** Two individuals were sitting next to each other and obtained food rewards from the two ends of the wooden board. As the task required no joint action, one individual could pull the string and move the board.

**Movie S3**

**Group service test condition.** The adult male in Pos. 0 pressed the handle, and as a result, the seesaw tilted. The adult female sitting in Pos. 1 obtained the food reward.

**Movie S4**

**Group service empty control condition.** The experimenter pretended to place food in Pos. 1. The individual did not press the handle within the maximum duration of the trial (2 minutes), suggesting an understanding of the contingency of the control condition.

**Movie S5**

**Blocked control condition.** The access to Pos. 1 was blocked using transparent plexiglass. Individuals noticed the food reward placed but could not obtain it. The individual did not press the handle within the maximum duration of the trial (2 minutes), suggesting an understanding of the contingency of the control condition.

**References**

1.	L. S. Pflüger, *et al.*, Twenty-three-year demographic history of the Affenberg Japanese macaques (Macaca fuscata), a translocated semi-free-ranging group in southern Austria. *Primates* **62**, 761–776 (2021).

2.	N. Nakagawa, “Intraspecific Differences in Social Structure of the Japanese Macaques: A Revival of Lost Legacy by Updated Knowledge and Perspective” in (2010), pp. 271–290.

3.	J. M. Burkart, C. van Schaik, Group service in macaques (Macaca fuscata), capuchins (Cebus apella) and marmosets (Callithrix jacchus): A comparative approach to identifying proactive prosocial motivations. *Journal of Comparative Psychology* **127**, 212–225 (2013).

4.	L. Horn, *et al.*, Sex-specific effects of cooperative breeding and colonial nesting on prosociality in corvids. *Elife* **9** (2020).

5.	J. J. M. Massen, C. Ritter, T. Bugnyar, Tolerance and reward equity predict cooperation in ravens (Corvus corax). *Scientific Reports* **5**, 15021 (2015).

6.	L. Horn, C. Scheer, T. Bugnyar, J. J. M. Massen, Proactive prosociality in a cooperatively breeding corvid, the azure-winged magpie ( *Cyanopica cyana* ). *Biology Letters* **12**, 20160649 (2016).

7.	A. Thornton, K. McAuliffe, Cognitive consequences of cooperative breeding? A critical appraisal. *Journal of Zoology* **295**, 12–22 (2015).

8.	E. C. Pielou, The measurement of diversity in different types of biological collections. *Journal of Theoretical Biology* **13**, 131–144 (1966).

9.	R Development Core Team, R Core Team (2020). R: A language and environment for statistical computing. R Foundation for Statistical Computing, Vienna, Austria. URL https://www.R-project.org/. *R Foundation for Statistical Computing* **2** (2019).

10.	D. Bates, M. Mächler, B. Bolker, S. Walker, Fitting Linear Mixed-Effects Models Using lme4. *Journal of Statistical Software* **67** (2015).

11.	T. Hothorn, A. Zeileis, Diagnostic Checking in Regression Relationships. *R News* **2** (2011).

12.	F. Hartig, DHARMa: Residual Diagnostics for Hierarchical Regression Models. *The Comprehensive R Archive Network* (2020).

13.	J. Fox, S. Weisberg, An {R} Companion to Applied Regression, Third Edition. *Thousand Oaks CA: Sage.* (2019).
